# Automated acquisition of knowledge beyond pathologists

**DOI:** 10.1101/539791

**Authors:** Yoichiro Yamamoto, Toyonori Tsuzuki, Jun Akatsuka, Masao Ueki, Hiromu Morikawa, Yasushi Numata, Taishi Takahara, Takuji Tsuyuki, Akira Shimizu, Ichiro Maeda, Shinichi Tsuchiya, Hiroyuki Kanno, Yukihiro Kondo, Manabu Fukumoto, Gen Tamiya, Naonori Ueda, Go Kimura

## Abstract

Deep learning algorithms have been successfully used in medical image classification and cancer detection. In the next stage, the technology of acquiring explainable knowledge from medical images is highly desired. Herein, fully automated acquisition of explainable features from annotation-free histopathological images is achieved via revealing statistical distortions in datasets by introducing the way of pathologists’ examination into a set of deep neural networks. As validation, we compared the prediction accuracy of prostate cancer recurrence using our algorithm-generated features with that of diagnosis by an expert pathologist using established criteria on 13,188 whole-mount pathology images. Our method found not only the findings established by humans but also features that have not been recognized so far, and showed higher accuracy than human in prognostic prediction. This study provides a new field to the deep learning approach as a novel tool for discovering uncharted knowledge, leading to effective treatments and drug discovery.

---

Trained on massive amounts of annotated data, deep learning algorithms have been successfully used in medical image classification and cancer detection. Esteva *et al.* successfully used a deep neural network to categorize fine-grained images of skin tumors, including malignant melanomas, at a dermatologist level^1^. Fauw *et al.* detected a range of sight-threatening retinal diseases as efficiently as an expert ophthalmologist, even on a clinically heterogeneous set of three-dimensional optical coherence tomographs (OCTs)^2^. Chilamkurthy *et al.* retrospectively collected a large annotated dataset of head computed tomography (CT) and evaluated the potential of deep learning algorithm to identify critical findings on CT images^3^. Bejnordi *et al.* evaluated the performance of deep learning algorithms submitted as part of a challenge competition and found that the performance of the high-ranking algorithm was comparable to that of pathologists in the detection of breast cancer metastases in histopathological tissue sections of lymph nodes^4^. Currently, machine learning-enhanced hardware is also being developed. Google has announced the development of an augmented reality microscope based on deep learning algorithms to assist pathologists^5^. However, automated acquiring explainable knowledge from medical images has not been uncharted.

Pathological examinations are used to provide definitive diagnoses and are one of the most reliable examinations in current cancer medicine^6^, but the diagnostic pathology knowledge and skills needed can only be acquired by completing a long fellowship program^7^. Although machine learning-driven histopathological image analysis^4, 8, 9^is an attractive tool to assist doctors, it faces two significant hurdles: the need for explainable analyses to gain clinical approval and the tremendous amount of information in histopathological images^8, 10^. Acquiring explainable knowledge from medical images is imperative for medicine. Furthermore, there are substantial size differences between histopathological images and other medical images^1-3, 11, 12^. A pathology slide contains large number of cells and the image consists of as many as tens of billion pixels^8^.

We aimed to develop a new method to acquire explainable features from annotation-free histopathological images and assessed the prediction accuracy of prostate cancer recurrence using our algorithm-generated features by comparison with that of human-established cancer criteria, the Gleason score by an expert in the diagnosis of prostate cancer.

## Results

First, we have developed a new method of generating key features that employs two different unsupervised deep neural networks (deep autoencoders^13, 14^) at different magnifications and weighted non-hierarchical clustering^15^ (**Fig. 1 and Supplementary Figures 1 and 2**). This takes histopathological images with over 10 billion pixel features and reduces them to only 100 clustered features with scores while retaining the images’ core information (**Fig. 2**). These clustered features can be effective for tasks such as predicting cancer recurrence, understanding the contributions of particular features and automatically annotating images. In the key feature generation dataset, short-term biochemical recurrence (BCR) cases were considered positive purely based on the recurrence time for patients (the recurrence period range: 1.7–14.4 months). No direct information regarding cancer images was provided to deep neural networks.

**Figure 1.**
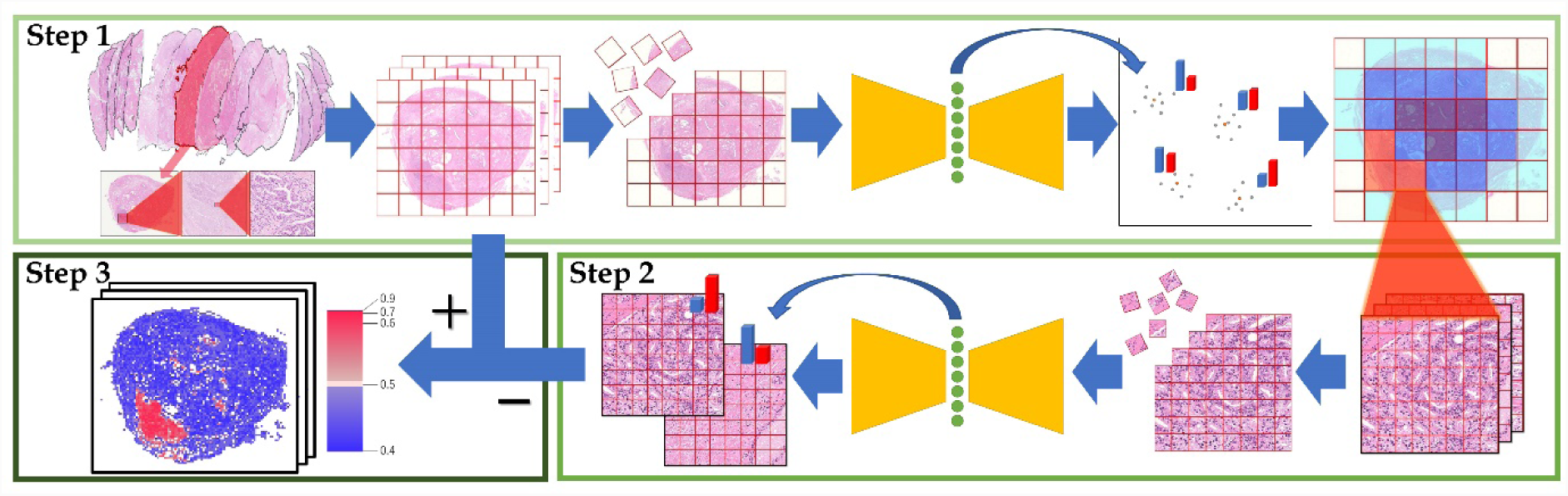
Key feature generation method. This approach was inspired by the way pathologists typically conduct diagnosis via step-by-step microscopic inspection. **Step 1**: First, we divide a low-magnification pathology images into smaller images, then perform dimensionality reduction using a deep autoencoder followed by weighted non-hierarchical clustering. This process reduces an image with over 10 billion-pixel features to only 100 clustered features with scores. (This step corresponds to the way pathologists search low-magnification images.) **Step 2**: Next, we analyse high-magnification images in order to reduce the number of misclassified low-magnification images. Again, we divide these into smaller images, before applying a second deep autoencoder and calculating average scores for the images. (This step corresponds to the way pathologists confirm their findings at a higher magnification.) **Step 3**: We remove images where the results of Steps 1 and 2 do not match. Finally, we use the total numbers of each type f clustered feature to make predictions. For example, to make cancer recurrence predictions, create human-understandable features or automatically annotate images.

**Figure 2.**
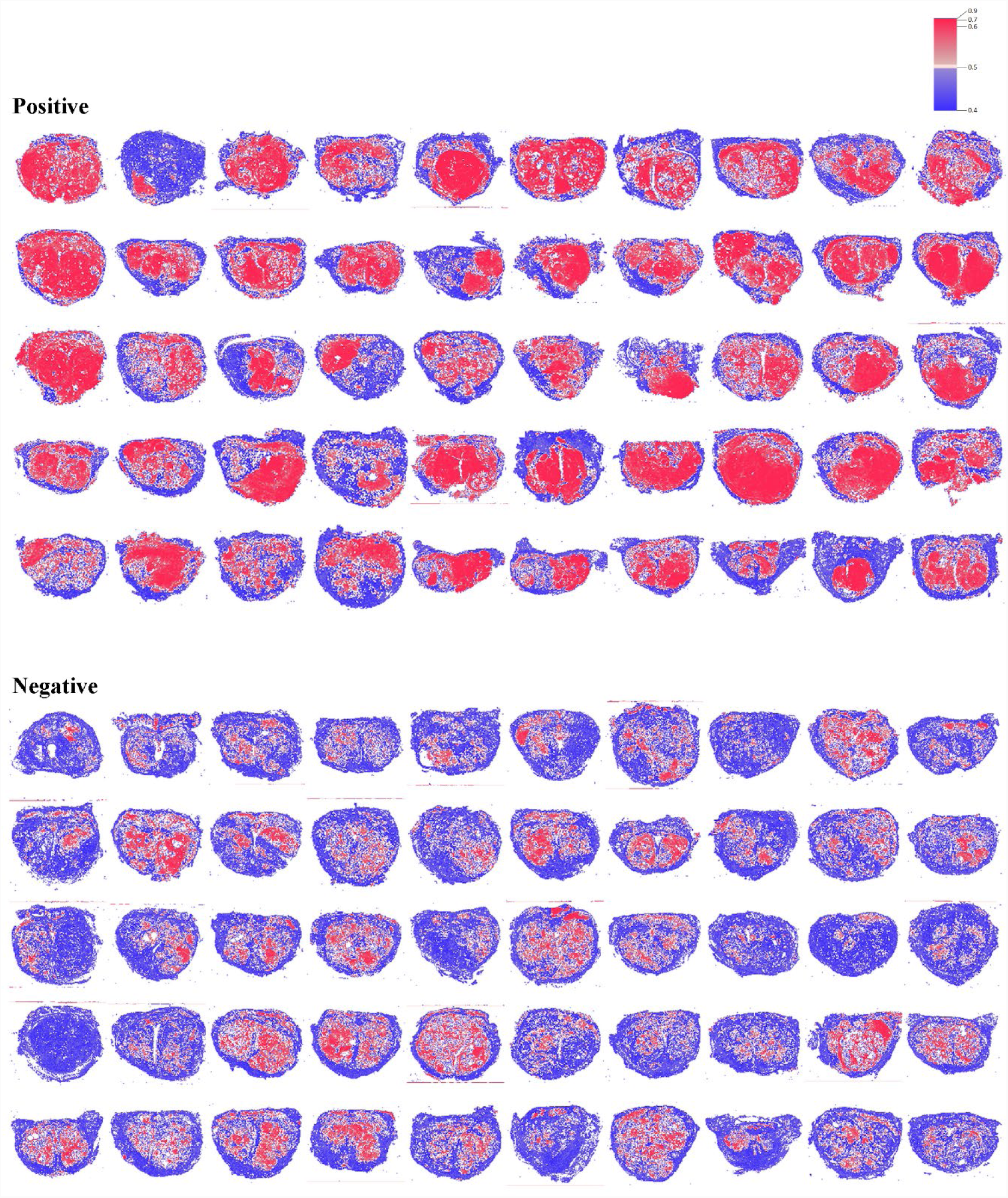
Examples of compressed images. Whole-mount pathology images with >10 billion-pixel features were reduced to only 100 clustered features, while retaining core image information. The color of each region indicates positive (red) and negative (blue) for characteristics detected.

Next, we validated the accuracy of cancer recurrence prediction using deep learning-generated features by comparing the predicted results with the Gleason score, one of the most crucial clinicopathological factors in the current prostate cancer practice^16^. The Gleason grading system defines five architectural growth patterns, which provides information on prostate cancer aggressiveness and facilitates patients’ appropriate care. As prostate cancer usually harbors two or more Gleason patterns, the sum of primary and secondary patterns yields the Gleason score. In this paper, all images’ Gleason score were evaluated by an expert genitourinary (GU) pathologist, T. Tsuzuki (the second author).

Our cohort included 1,007 patients with prostate cancers who received a radical prostatectomy, with a total of 13,188 whole-mount pathology slides. We excluded 115 cases involving neoadjuvant therapy and 7 cases involving adjuvant therapy as well as 43 cases who could not be followed up within 1 year because of hospital transfer or death due to other causes, thus leaving 842 cases for analysis. **Table 1** summarizes the clinical characteristics of the study cohort. Cancer was more likely to recur in patients with higher prostate-specific antigen (PSA) levels (*P* < 0.001). It was more likely to recur in patients with a higher Gleason score (≥8) than in patients with a lower Gleason score (<8). Similar patterns were observed in 1-year and 5-year recurrence rates. No significant differences existed in the average age, height, weight, or prostate weight between patients in whom cancer recurred and those in whom it did not. We categorized the data for 842 patients into the following two sets: 100 patients (100 whole-mount pathology images) were used to generate key features using the deep neural networks; and 742 (9,816 images) were used to perform the BCR predictions using these features. We applied lasso^17^and ridge^18^regression analyses and a support vector machine (SVM)^19^to the clustered features to predict the BCR of prostate cancer. We evaluated the areas under the receiver operating characteristic curves (AUCs) with a 95% confidence interval (CI) and receiver operating characteristic (ROC) curves^20, 21^. **Table 2** and **Fig. 3** present the AUCs and ROC curves of BCR predicted using the deep learning-generated features and we compared these values to the Gleason score. The AUC for BCR predictions by the deep neural networks within 1 year was 0.82 (95% CI: 0.766–0.873), while the Gleason score was 0.744 (95% CI: 0.672– 0.816). Interestingly, combining both methods produced a more accurate BCR prediction [AUC, 0.842 (95% CI: 0.788–0.896)] than either method alone. Likewise, the 5-year prediction accuracies were 0.721 (95% CI: 0.672–0.769; deep neural networks), 0.695 (95% CI: 0.639– 0.75; Gleason score), and 0.758 (95% CI: 0.71–0.806; combined).

**Table 1.**
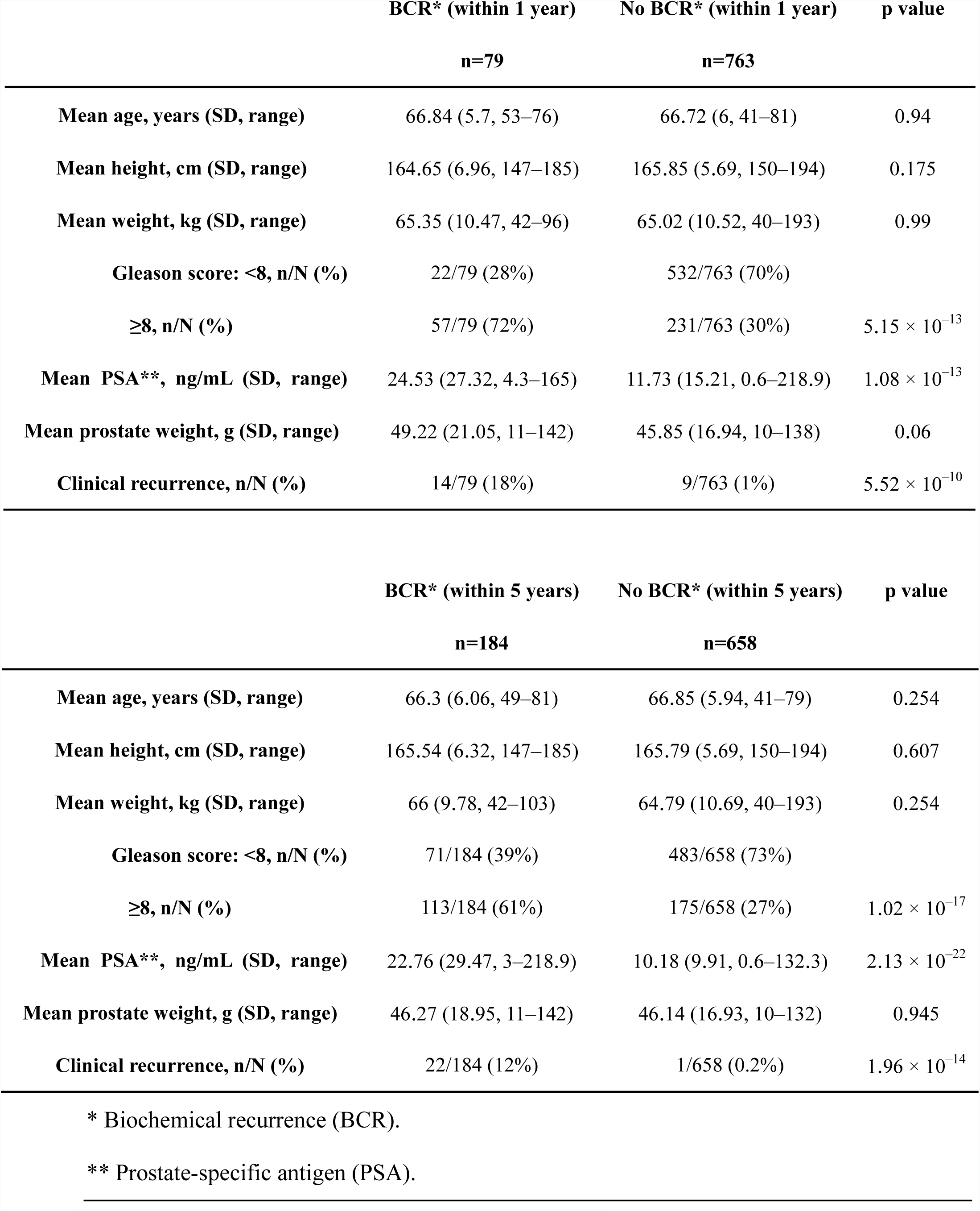
The clinical characteristics of the cohort

|  | BCR* (within 1 year) | No BCR* (within 1 year) | p value |
| --- | --- | --- | --- |
|  | n=79 | n=763 |  |
| Mean age, years (SD, range) | 66.84 (5.7, 53–76) | 66.72 (6, 41–81) | 0.94 |
| Mean height, cm (SD, range) | 164.65 (6.96, 147–185) | 165.85 (5.69, 150–194) | 0.175 |
| Mean weight, kg (SD, range) | 65.35 (10.47, 42–96) | 65.02 (10.52, 40–193) | 0.99 |
| Gleason score: <8, n/N (%) | 22/79 (28%) | 532/763 (70%) |  |
| ≥8, n/N (%) | 57/79 (72%) | 231/763 (30%) | $5.15 \times 10^{-13}$ |
| Mean PSA**, ng/mL (SD, range) | 24.53 (27.32, 4.3–165) | 11.73 (15.21, 0.6–218.9) | $1.08 \times 10^{-13}$ |
| Mean prostate weight, g (SD, range) | 49.22 (21.05, 11–142) | 45.85 (16.94, 10–138) | 0.06 |
| Clinical recurrence, n/N (%) | 14/79 (18%) | 9/763 (1%) | $5.52 \times 10^{-10}$ |

**Table 1. The clinical characteristics of the cohort**
|  | BCR* (within 5 years) | No BCR* (within 5 years) | p value |
| --- | --- | --- | --- |
|  | n=184 | n=658 |  |
| Mean age, years (SD, range) | 66.3 (6.06, 49–81) | 66.85 (5.94, 41–79) | 0.254 |
| Mean height, cm (SD, range) | 165.54 (6.32, 147–185) | 165.79 (5.69, 150–194) | 0.607 |
| Mean weight, kg (SD, range) | 66 (9.78, 42–103) | 64.79 (10.69, 40–193) | 0.254 |
| Gleason score: <8, n/N (%) | 71/184 (39%) | 483/658 (73%) |  |
| ≥8, n/N (%) | 113/184 (61%) | 175/658 (27%) | $1.02 \times 10^{-17}$ |
| Mean PSA**, ng/mL (SD, range) | 22.76 (29.47, 3–218.9) | 10.18 (9.91, 0.6–132.3) | $2.13 \times 10^{-22}$ |
| Mean prostate weight, g (SD, range) | 46.27 (18.95, 11–142) | 46.14 (16.93, 10–132) | 0.945 |
| Clinical recurrence, n/N (%) | 22/184 (12%) | 1/658 (0.2%) | $1.96 \times 10^{-14}$ |

\* Biochemical recurrence (BCR).
\*\* Prostate-specific antigen (PSA).

**Table 2.** AUC comparison

|  | BCR* (within 1 year) | BCR* (within 5 years) |
| --- | --- | --- |
| Gleason score (pathologist) | **0.744 [95% CI 0.672–0.816]** | **0.695 [95% CI 0.639–0.75]** |
| Ridge (automated) | 0.801 [95% CI 0.748–0.854] | 0.696 [95% CI 0.647–0.744] |
| Lasso (automated) | 0.804 [95% CI 0.749–0.86] | 0.684 [95% CI 0.634–0.734] |
| SVM (automated) | **0.82 [95% CI 0.766–0.873]** | **0.721 [95% CI 0.672–0.769]** |
| Ridge + Gleason score | 0.824 [95% CI 0.77–0.878] | 0.732 [95% CI 0.684–0.78] |
| Lasso + Gleason score | 0.83 [95% CI 0.772–0.888] | 0.735 [95% CI 0.688–0.783] |
| SVM + Gleason score | **0.842 [95% CI 0.788–0.896]** | **0.758 [95% CI 0.71–0.806]** |

The reported values are averages with 95% confidence interval. The bold values are the best accuracy of lasso, ridge and SVM.
\*Biochemical recurrence (BCR).

**Figure 3.**
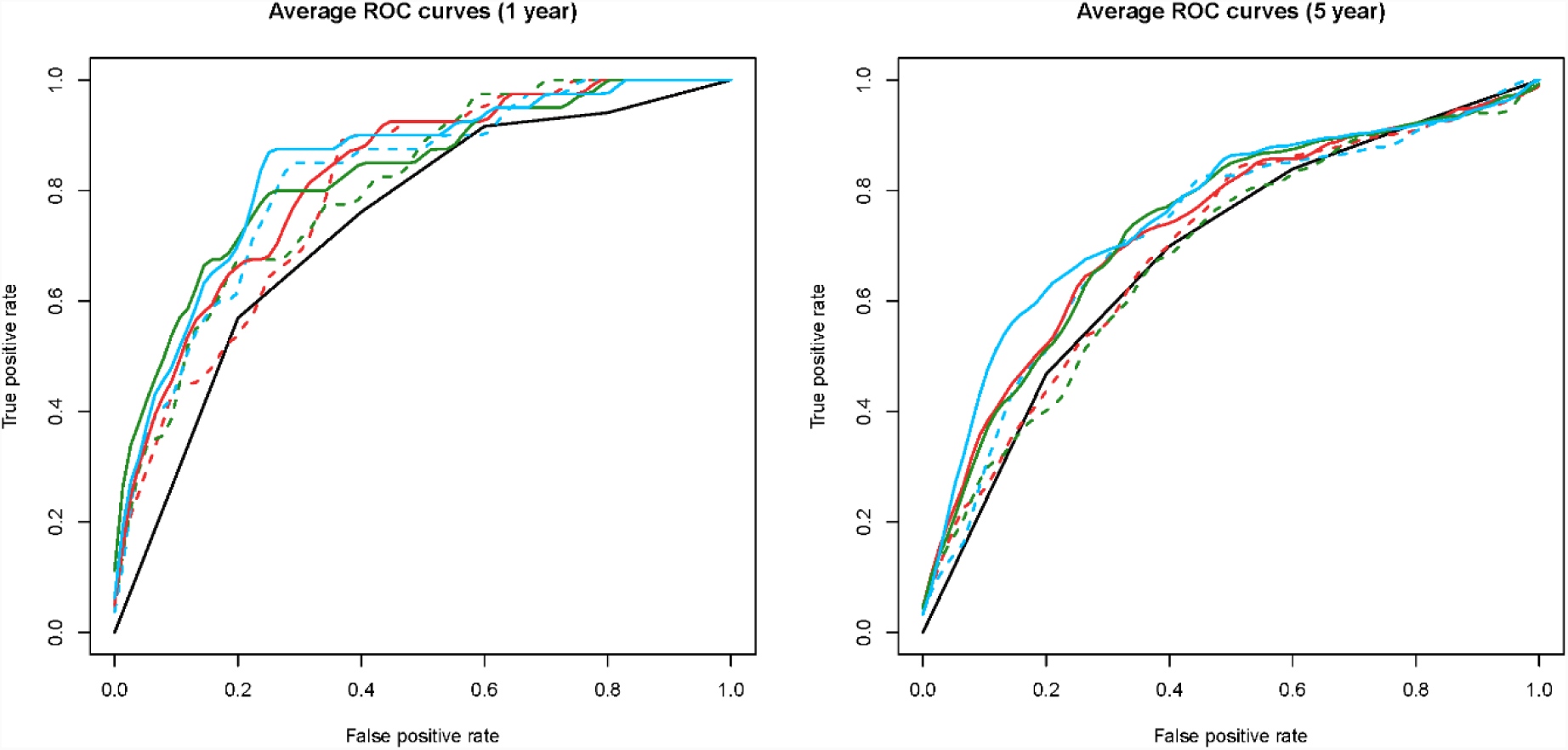
Receiver operating characteristic (ROC) curves for the biochemical recurrence (BCR) prediction. Average ROC curves for the BCR prediction within one year (left) and BCR prediction within five years (right). The Gleason score (black solid line), Ridge (red dot line), Lasso (green dot line), support vector machine (SVM; blue dot line), Ridge + Gleason score (red solid line), Lasso + Gleason score (green solid line), SVM + Gleason score (blue solid line).

Then, we selected the images that were closest to each cluster’s centroid as being representative of the clustered features (**Fig. 4**). The expert GU pathologist (T. Tsuzuki) analyzed these images to search for pathological meanings (**Table 3**). In summary, the pathologist found that the deep neural networks appeared to have mastered the basic concept of the Gleason score, fully automatically, generating explainable key features that could be understood by pathologists. Furthermore, the deep neural networks identified the features of stroma in the noncancerous area as a prognostic factor, which typically have not been evaluated in prostate histopathological images. **Fig. 5** and **supplementary videos 1–2** show feature maps for a whole-mount pathology image as well as cell-level information about images; the predicted high-grade cancer regions are shaded in red, whereas normal ducts/low-grade cancer regions are shaded in blue.

**Table 3.**
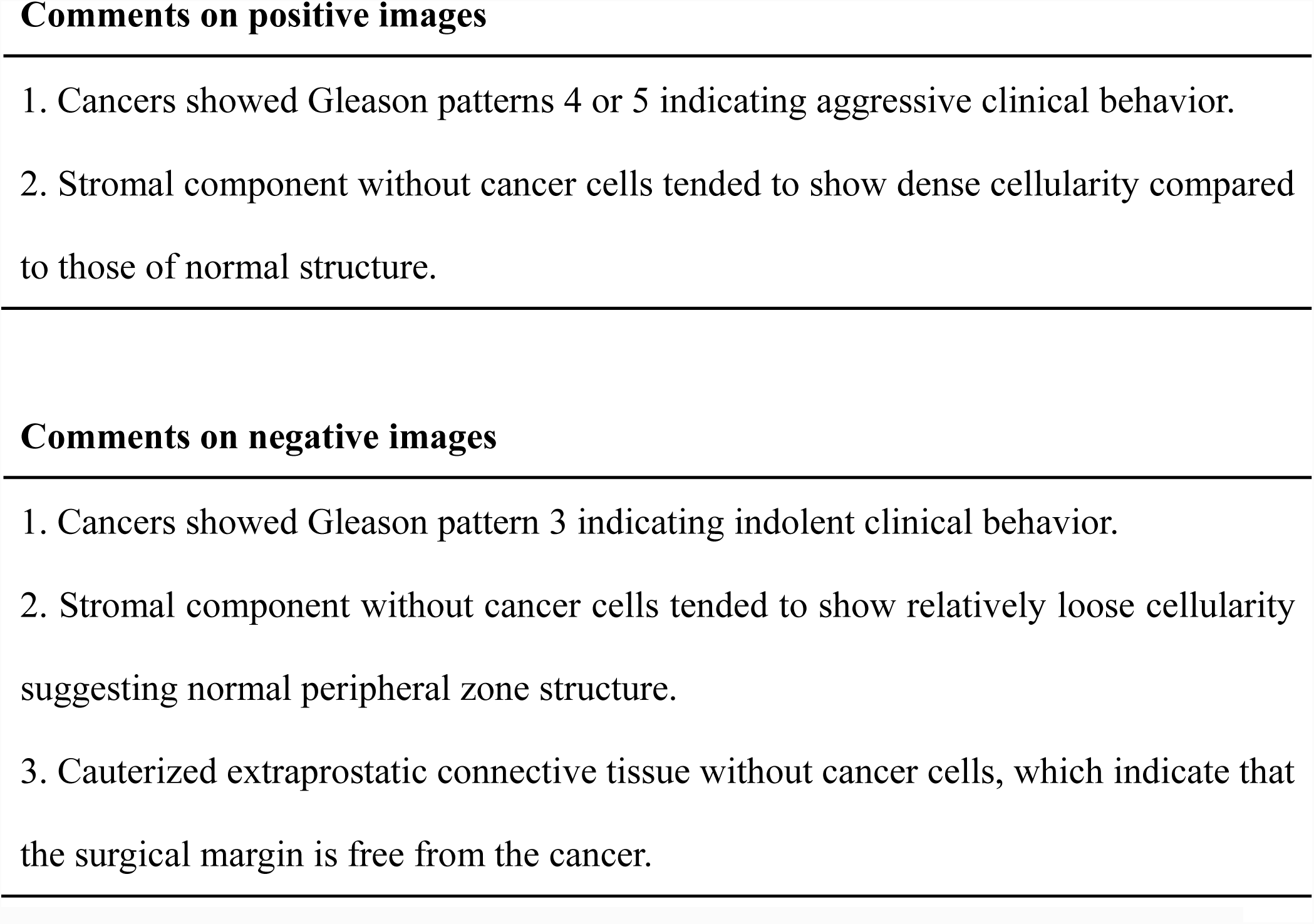
Expert genitourinary (GU) pathologist’s comments on figure 4.

**Figure 4.**
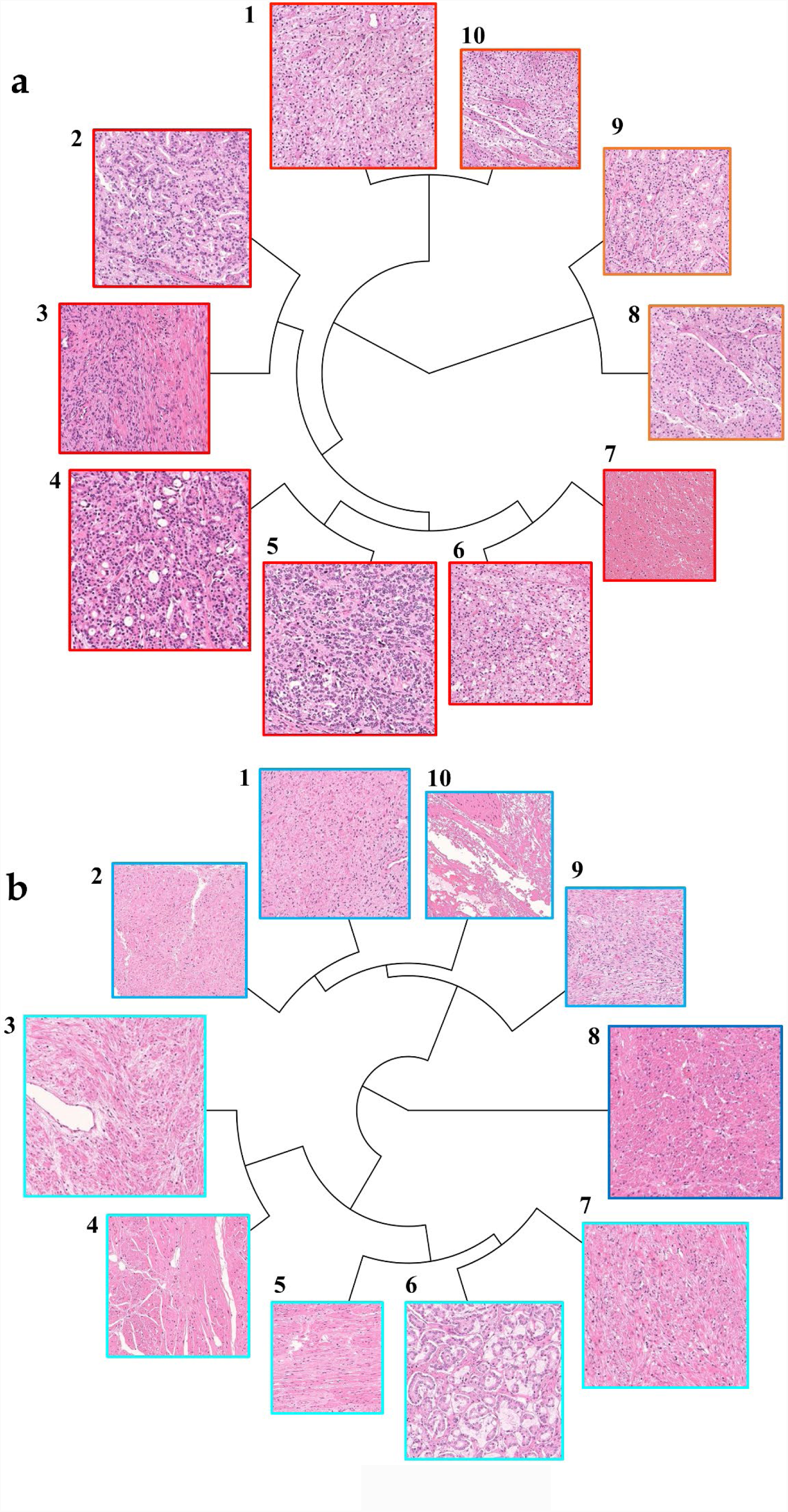
Representative images of key features. The top 10 images are closest to the centroids of the 100 clusters, with higher-ranking images being larger, in the (a) biochemical recurrence (BCR) group and (b) no BCR group (see also Table 3). (a) 1,2,4,5,6,8,9,10: Cancers equivalent to Gleason patterns 4 or 5, which usually indicate aggressive clinical behavior. 3: Dense stromal components without cancer cells. 7: Hemorrhage. (b) 6: Cancers equivalent to Gleason pattern 3, which usually indicates benign clinical behavior. 1,2,3,4,5,7,8,9: Loose stromal components without cancer cells. 10: Surgical margin without cancer cells.

**Figure 5.**
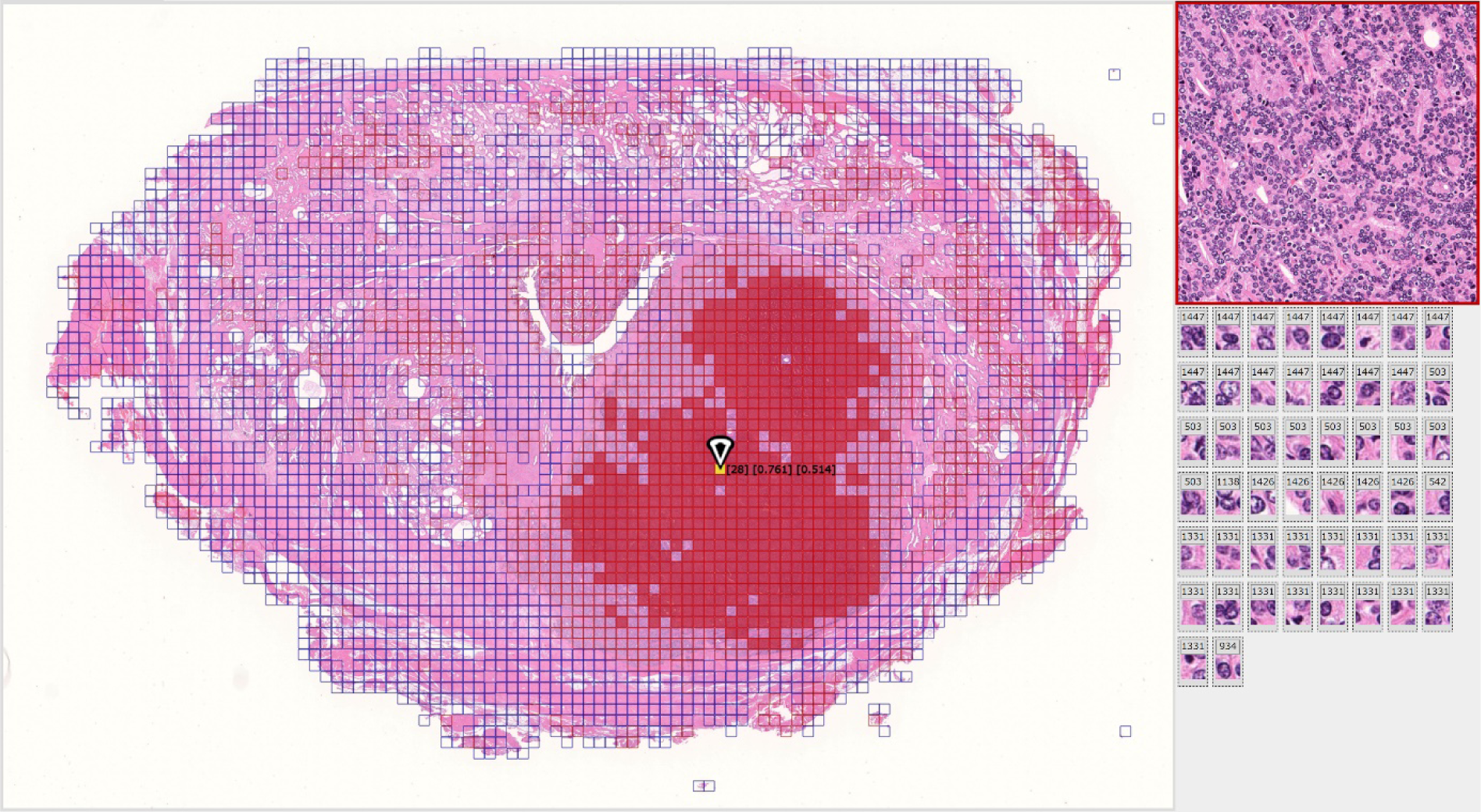
Automatically annotated whole-mount pathology image based on key features and cell-level information. Our method directly generates key features based on the whole image, without requiring a region selection step. Using the key features and cell-level information found by the deep neural networks we automatically annotated a whole-mount pathology image. Here we show an automatically annotated whole-mount pathology image (left), as well as a low-magnification image of the yellow region (upper right) and the associated high-magnification images (lower right). The regions with impact scores above and below 0.5 in Step 1 are shaded in red and blue, respectively. The indicated cell shows [number of clusters] [impact score, Step 1] [impact score, Step 2] (see ‘Key feature generation method’ in the methods section).

## Discussion

We achieved fully automated acquisition of explainable features from histopathological images in the raw. Our method found not only the human-established findings but also previously-unrecognized pathological features, resulting in higher prediction accuracy of cancer recurrence than that of diagnosis performed by an expert pathologist using human-established cancer criteria, the Gleason score.

The Gleason score^22^ is a unique pathological grading system, purely based on architectural disorders, without considering cytological atypia. In this study, none of the cancer cells in the images identified by the deep neural networks as representative of high-grade cancer showed severe nuclear atypia or prominent nucleoli. Our results of the deep neural networks indicate that the central ideas behind Gleason’s grading system are sound.

The most accurate BCR predictions was produced by combining the deep learning-generated features and Gleason score, possibly because the automatically derived features included factors different from those used for the Gleason score, such as the surgical margin status. Various and complex factors are believed to be associated with BCR^23, 24^. Interestingly, representative images of the features nominated by the deep neural networks comprised not only the human-established findings but also previously unspotlighted or neglected features of stroma at the noncancerous area. These findings indicate that the deep neural networks could explore unique features that could be underestimated or overlooked by a human.

In this study, the deep neural networks identified comprehensible key features from scratch. Silver *et al*. reported that the AlphaGo Zero^25^ program, which is solely based on reinforcement learning without any human knowledge inputs, could defeat their previous AlphaGo^26^ program, which conducted supervised learning using human expert moves. In this study, we demonstrated another algorithm that performs well, is based on deep autoencoders^13, 14^, and does not need human knowledge. Hopefully, this algorithm will provide a novel tool for discovering new findings. In addition, our method can be applied to non-verbal information, such as that derived from the subjective experience of experts, as long as it is used to classify images. For example, data from patients with similar symptoms but unknown causes could be used to discover the key underlying factors, resulting in more effective treatments and the development of new medicines. We anticipate that our method will lead to the new design of clinical trials using deep learning and therapeutic strategies and will help reduce the workloads of busy physicians^27^.

This study has some limitations. Our results are limited in that we did not perform validation in multiple centers. A clinical trial is required to determine whether our method is universally effective for improving the prediction accuracy and patient care in different areas. Nonetheless, our cohort was sufficiently large and provides reliable and robust results in one facility. Furthermore, we present detailed flowcharts and methods in this paper, and all processes are sufficiently described to enable independent replication, warranting evaluation using a larger global patient cohort.

Human and computer analyses have different strengths. Our deep learning approach analyses huge medical images broadly and without oversights or bias; a human pathologist analyses the disease more accurately and with a greater focus on medical importance. Each approach can, therefore, complement the other. Medicine aims to save patients, and both medical doctors and artificial intelligence (AI) systems can contribute to this goal. The more effectively and deeply medical experts can utilize AI systems, the more patients will benefit. Increasing collaboration between medical experts and informaticians will surely improve outcomes for patients.

## Methods

### Subjects and Ethics

This hospital-based cohort comprised all patients with prostate cancers who received a radical prostatectomy from April 2000 to December 2016 at the Nippon Medical School Hospital (*N* = 1,007). We collected whole-mount pathology slides and clinical data for all patients in this cohort. Of note, no patients were enrolled on clinical trials of radical prostatectomy. All patients were followed and checked for the BCR every 3 months postoperatively; the median follow-up duration was 72.8 months. We defined the BCR following radical prostatectomy based on the European Association of Urology guidelines of increasing PSA levels >0.2 ng/mL^28^. We excluded 115 cases involving neoadjuvant therapy and 7 cases involving adjuvant therapy as well as 43 cases who could not be followed up within 1 year because of hospital transfer or death due to other causes, thus leaving 842 cases for analysis. This research was approved by the Institutional Review Boards of the Nippon Medical School Hospital (reference 28-11-663) and RIKEN (reference Wako3 29-14), Japan.

### Datasets

We categorized the data for 842 patients into the following two sets: 100 patients (100 whole-mount pathology images) were used to generate key features using the deep neural networks; and 742 (9,816 images) were used to perform BCR predictions using these features. We carefully ensured that no direct information regarding cancer concepts was provided to deep neural networks. In addition, histopathological images were not checked or annotated by pathologists before key feature generation was performed by the deep neural networks. In the key feature generation dataset, short-term BCR cases were considered positive purely based on the recurrence time for patients (the recurrence period range: 1.7–14.4 months). To avoid bias, we also used the same surgery year distribution to select negative cases. Of note, images that extended beyond the edge of the cover glass were not used for key feature generation. During the key feature generation process, we simply selected the largest available image in each patient, without checking whether any cancer was included.

### Statistical analysis

We compared the characteristics of patients whose cancer did or did not recur within 1 and 5 years postoperatively using the Fisher’s exact test for categorical data and the Wilcoxon rank-sum test for continuous data (**Table 1**). All tests were two-tailed and were considered statistically significant if *P* < 0.05. All statistical analyses were performed using R, version 3.4.4.

### Preparation of whole-mount pathology images

Whole prostates were fixed in 10% formalin and embedded in paraffin. All samples were sectioned at a thickness of 3 μm and stained with hematoxylin and eosin (H&E). All H&E-stained slides were scanned by a whole-slide imaging scanner (Hamamatsu NanoZoomer S60 Slide Scanner) with a 20× objective lens and were stored on a secure computer.

### Histological grading

We classified prostate cancer histologically based on the International Society of Urological Pathologists (ISUP) classification criteria^16^. All slides were initially reviewed independently by two board-certified pathologists, and our conclusions were confirmed by an expert GU pathologist (T. Tsuzuki) without using clinical data, including the BCR.

### Key feature generation method

The proposed method does not require human annotation for image classification and reveals statistical distortions in image datasets by employing multiple deep autoencoders at different magnifications and weighted nonhierarchical clustering. **Supplementary Figures 1 and 2** provide detailed algorithm flowcharts and descriptions of the autoencoder networks. Most previous methods include a region selection step, for example to extract or annotate the region of interest. In contrast, our method derives the key features directly from the whole image, without requiring such a step. It can be regarded as a type of dimensional reduction, and was inspired by the step-by-step microscopic inspection process pathologists typically use for diagnosis.

**Step 1**: We generated the key features from 100 whole-mount pathology images (100 cases), taken at low magnification (25x). We divided each image (considered as an image data vector ***S***_*i*_) into a set of small 128×128-pixel images ***S***_*i,j*_ using NDP.convert software (Hamamatsu Photonics K.K., version 2.0.7.0). We then applied a deep autoencoder we had developed for pathology images (**Supplementary Figure 2**) to each small image, clustering the 2048 intermediate-layer features to form 100 features by *k*-means clustering. Clusters that included white background areas without tissue were automatically removed. Next, we found the centroid of each cluster, and calculated a score *u*_*i,j,k*_ for each feature based on the distance from each centroid *d*_*i,j.k*_. Here, we applied the simplest possible scoring method, as follows:

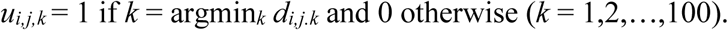

Defining the total number of small images belonging to the positive and negative groups and *n*_positive_ and *n*_negative_, respectively, we defined the positive and negative degrees *r*_positive, *k*_ and *r*_negative, *k*_ for the *k*th feature as

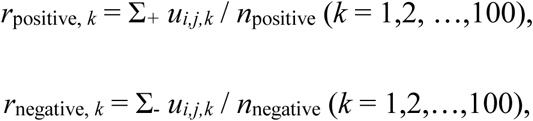

where the sums Σ_*+*_ and Σ_-_ are over all *i,j* pairs such that image ***S***_*i,j*_ belonged to the positive and negative groups, respectively. Finally, we defined the impact score *I*_*k*_ for the *k*th feature and the impact score *I*_*i,j*_ of image ***S***_*i,j*_ for this step as

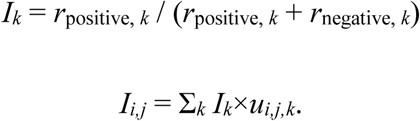

Step 1 corresponds to the way pathologists search low-magnification images.

**Step 2**: Next, high-magnification (200x) images were analysed to reduce the number of misclassified low-magnification images. Here, 1024×1024-pixel images for each of the small images in Step 1, considered as image data vectors ***S****′*_*i,j*_, were divided into small 28×28 pixel images ***S****′*_*i,j,j′*_. A second deep autoencoder (**Supplementary Figure 2**) was then applied to each of these smaller images. The 1,568 intermediate-layer features ***v****′*_*i,j,j′*_ were given scores *u′*_*i,j,j′,k′*_ based on the intensity values of each node. Again, we used the following simple scoring method:

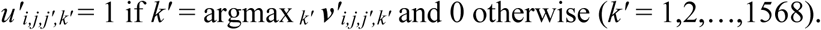

Defining the total number of small images belonging to the positive and negative groups as *n′*_positive_ and *n′*_negative_, we defined the positive and negative degrees *r′*_positive, *k′*_ and *r′*_negative, *k′*_ for the *k′*th feature as

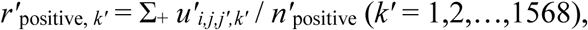

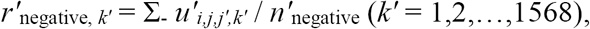

where the sums Σ_*+*_ and Σ_-_, analogously to those in Step 1, are over all *i,j,j′* such that the image ***S****′*_*i,j,j′*_ belonged to the positive and negative groups, respectively. For this step, we defined the impact score *I′*_*i,j*_ as

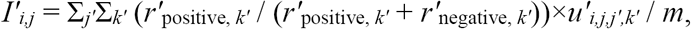

where *m* denotes the total number of small images ***S****′*_*i,j,j′*_ used for ***S***_*i,j*_.

Step 2 corresponds to the way pathologists confirm their findings at higher magnification.

**Step 3**: Images that were frequently in the positive and negative groups had impact scores above and below 0.5, respectively, so we defined images with impact scores above and below 0.5 as having positive and negative characteristics, respectively. We then removed images whose characters, based on the impact scores in Steps 1 and 2, did not match. Finally, we used the total numbers of each clustered feature type for the subsequent predictions.

### AUC comparison

To evaluate our approach, we predicted cancer recurrence using 9,816 whole-mount pathology images (742 cases), excluding 100 cases that were used for key feature generation. In particular, we assessed the potential of the 100 clustered features to predict the recurrence of cancer within 1 or 5 years postoperatively using Lasso^17^and Ridge^18^regression and a support vector machine (SVM)^19^, all popular methods for building prediction models. In addition, we created prediction models based on the application of logistic regression to an ISUP grade group assessed on the basis of the Gleason score and similarly created models combining the 100 clustered features with the grade. If multiple images were available for a given patient, we averaged each feature over all the images. To address the fact that the feature values were not evenly distributed amongst patients where cancer did and did not recur, we multiplied each feature value by 1 + | *I*_*k*_ –0.5| (see ‘Key feature generation method’ in the methods section), which augmented the predictive power of the models. We used 10-fold cross-validation^29, 30^to test the prediction models, randomly dividing the whole sample set in a 1:9 ratio, using one part for testing and the other nine parts for training. For each testing/training split, we used the AUC metric to assess the performance of trained prediction models on the test data^20, 21^. We used R for the analysis, using the glmnet package (version 2.0.16) for Ridge and Lasso regression, the e1071 package (version 1.7.0) for the SVM, and the cvAUC package to evaluate the AUC with a CI.

## Data availability

The clinical data used for the training and test sets were collected at the Nippon Medical School Hospital. This work and the collection of data was approved by the Institutional Review Boards of the Nippon Medical School Hospital. They are not publicly available, and restrictions apply to their use.

**Supplementary Figure legends**

**Supplementary Figure 1** | Algorithm Flowcharts.

**Supplementary Figure 2** | Networks of deep autoencoders.

**Supplementary Videos 1-2**

Video1.mp4

Video2.mp4

## Acknowledgment

This study was conducted by the RIKEN AIP Deep Learning Environment (RAIDEN) supercomputer system for the computations. We thank the RAIDEN-supporting members at the RIKEN AIP center. We also thank Prof. Takeo Kanade for his insight. This research was supported by the ICT Infrastructure for the Establishment and Implementation of Artificial Intelligence for Clinical and Medical Research of the Japan Agency for Medical Research and development, AMED, and the Centre for Advanced Intelligence Project, RIKEN. We are currently applying for patents on the method presented in this paper.

## Author Contributions

Y.Y. designed this study, invented the method, programmed the machine learning system, analysed the data and wrote the manuscript. T.Tsuzuki performed pathological diagnoses, evaluated the Gleason score of all slides and helped with both writing the manuscript and discussion. J.A. digitised the histopathological slides, constructed the dataset and participated in discussions. M.U. conducted statistical analyses of the dataset and AUC comparisons. H.M. programmed the machine learning system and helped with data analyses. Y.N. helped with the programming of the machine learning system and analysed the data. T.Takahara and T.Tsuyuki performed pathological diagnoses and evaluated the Gleason score. A.S. and his laboratory members made whole-mount histopathology slides and helped with pathological discussion and diagnosis. I.M., S.T. and H.K., helped with pathological discussion and diagnosis. Y.K. helped with dataset construction and clinical discussion. F.M. helped with writing the manuscript and discussion. G.T. helped with discussion and statistical analysis. N.U. helped with discussion and supervised the study. G.K. designed the study, constructed the dataset, helped with discussion and supervised the study.

## Author Information

The authors declare no competing financial interests. Correspondence and requests should be addressed to Y.Y. or G.K.

## References

1 Esteva, A. et al. Dermatologist-level classification of skin cancer with deep neural networks. Nature 542, 115–118, doi:10.1038/nature21056 (2017).

2 De Fauw, J. et al. Clinically applicable deep learning for diagnosis and referral in retinal disease. Nat Med, doi:10.1038/s41591-018-0107-6 (2018).

3 Chilamkurthy, S. et al. Deep learning algorithms for detection of critical findings in head CT scans: a retrospective study. Lancet, doi:10.1016/S0140-6736(18)31645-3 (2018).

4 Ehteshami Bejnordi, B. et al. Diagnostic Assessment of Deep Learning Algorithms for Detection of Lymph Node Metastases in Women With Breast Cancer. JAMA 318, 2199–2210, doi:10.1001/jama.2017.14585 (2017).

5 Stumpe, M. An Augmented Reality Microscope for Cancer Detection, https://ai.googleblog.com/2018/04/an-augmented-reality-microscope.html (2018).

6 Connolly JL S. S., Wang HH. Role of the Surgical Pathologist in the Diagnosis and Management of the Cancer Patient. 6th edition (BC Decker, 2003).

7 Barger, L. K. et al. Extended work shifts and the risk of motor vehicle crashes among interns. N Engl J Med 352, 125–134, doi:10.1056/NEJMoa041401 (2005).

8 Daisuke Komura, S. I. Machine Learning Methods for Histopathological Image Analysis. Computational and Structural Biotechnology Journal 16, 34–42 (2018).

9 Yamamoto, Y. et al. Quantitative diagnosis of breast tumors by morphometric classification of microenvironmental myoepithelial cells using a machine learning approach. Sci Rep 7, 46732, doi:10.1038/srep46732 (2017).

10 Gurcan, M. N. et al. Histopathological image analysis: a review. IEEE Rev Biomed Eng 2, 147–171, doi:10.1109/RBME.2009.2034865 (2009).

11 Lakhani, P. & Sundaram, B. Deep Learning at Chest Radiography: Automated Classification of Pulmonary Tuberculosis by Using Convolutional Neural Networks. Radiology 284, 574–582, doi:10.1148/radiol.2017162326 (2017).

12 Kim, K. et al. Performance of the deep convolutional neural network based magnetic resonance image scoring algorithm for differentiating between tuberculous and pyogenic spondylitis. Sci Rep 8, 13124, doi:10.1038/s41598-018-31486-3 (2018).

13 Rumelhart, D. E., Hinton, G. E. & Williams, R. J. Learning internal representations by error propagation. Parallel Distributed Processing. Vol 1: Foundations. (MIT Press, Cambridge, MA, 1986).

14 Hinton, G. E. & Salakhutdinov, R. R. Reducing the dimensionality of data with neural networks. Science 313, 504–507, doi:10.1126/science.1127647 (2006).

15 Arthur, D. & Vassilvitskii, S. Society for Industrial and Applied Mathematics Philadelphia, PA, USA. 1027–1035 (2007).

16 Epstein, J. I. et al. The 2014 International Society of Urological Pathology (ISUP) Consensus Conference on Gleason Grading of Prostatic Carcinoma: Definition of Grading Patterns and Proposal for a New Grading System. Am J Surg Pathol 40, 244–252, doi:10.1097/PAS.0000000000000530 (2016).

17 Tibshirani, R. Regression shrinkage and selection via the Lasso. J Roy Stat Soc B Met 58, 267–288 (1996).

18 Hoerl, A. E. & Kennard, R. W. Ridge Regression-Biased Estimation for Nonorthogonal Problems. Technometrics 12, 55–67 (1970).

19 Vapnik, V. Statistical Learning Theory. (John Wiley and Sons, 1998).

20 Pirracchio, R. et al. Mortality prediction in intensive care units with the Super ICU Learner Algorithm (SICULA): a population-based study. Lancet Respir Med 3, 42–52, doi:10.1016/S2213-2600(14)70239-5 (2015).

21 LeDell, E., Petersen, M. & van der Laan, M. Computationally efficient confidence intervals for cross-validated area under the ROC curve estimates. Electron J Stat 9, 1583- 1607, doi:10.1214/15-EJS1035 (2015).

22 Phillips, J. L. & Sinha, A. A. Patterns, art, and context: Donald Floyd Gleason and the development of the Gleason grading system. Urology 74, 497–503, doi:10.1016/j.urology.2009.01.012 (2009).

23 Tsuzuki, T. Intraductal carcinoma of the prostate: a comprehensive and updated review. Int J Urol 22, 140–145, doi:10.1111/iju.12657 (2015).

24 Kato, M. et al. Integrating tertiary Gleason pattern 5 into the ISUP grading system improves prediction of biochemical recurrence in radical prostatectomy patients. Mod Pathol, doi:10.1038/s41379-018-0121-8 (2018).

25 Silver, D. et al. Mastering the game of Go without human knowledge. Nature 550, 354–359, doi:10.1038/nature24270 (2017).

26 Silver, D. et al. Mastering the game of Go with deep neural networks and tree search. Nature 529, 484–489, doi:10.1038/nature16961 (2016).

27 Robboy, S. J. et al. Pathologist workforce in the United States: I. Development of a predictive model to examine factors influencing supply. Arch Pathol Lab Med 137, 1723–1732, doi:10.5858/arpa.2013-0200-OA (2013).

28 Cornford, P. et al. EAU-ESTRO-SIOG Guidelines on Prostate Cancer. Part II: Treatment of Relapsing, Metastatic, and Castration-Resistant Prostate Cancer. Eur Urol 71, 630–642, doi:10.1016/j.eururo.2016.08.002 (2017).

29 Stone, M. Cross-Validatory Choice and Assessment of Statistical Predictions. J R Stat Soc B 36, 111–147 (1974).

30 Hastie, T., Tibshirani, R. & Friedman, J. H. The elements of statistical learning : data mining, inference, and prediction. 2nd edition (Springer, 2009).

